# Desmosomal dualism: the core is stable while plakophilin is dynamic

**DOI:** 10.1101/2021.03.02.433631

**Authors:** Judith B Fülle, Henri Huppert, David Liebl, Jaron Liu, Rogerio Alves de Almeida, Bian Yanes, Graham D Wright, E Birgitte Lane, David R Garrod, Christoph Ballestrem

## Abstract

Desmosomes, strong cell-cell junctions of epithelia and cardiac muscle, link intermediate filaments to cell membranes and mechanically integrate cells across tissues, dissipating mechanical stress. They comprise 5 major protein classes - desmocollins and desmogleins (the desmosomal cadherins), plakoglobin, plakophilins and desmoplakin - whose individual contribution to the structure and turnover of desmosomes is poorly understood. Using live-cell imaging together with FRAP and FLAP we show that desmosomes consist of two contrasting protein fractions or modules: a very stable desmosomal core of desmosomal cadherins and plakoglobin, and a highly mobile plakophilin. As desmosomes mature from calcium-dependence to calcium-independent hyper-adhesion, core stability increases, but Pkp2a remains highly mobile. Desmoplakin is initially mobile but stabilises with hyper-adhesion. We show that desmosome down-regulation during growth factor-induced cell scattering proceeds by internalisation of whole desmosomes, which still retain a stable core and highly mobile Pkp2a. This molecular mobility of Pkp2a suggests a transient and probably regulatory role for Pkp2a in the desmosome.

## Introduction

Vertebrates have evolved several multimolecular systems to provide the cell cohesion and resilience necessary to withstand mechanical stress and deformation without a protective exoskeleton. Epithelia, the functional interfaces around, within and between body organs, have evolved complex multiprotein desmosome cell-cell junctions. Desmosomes connect to cytoplasmic intermediate filament networks to form a mechanically resilient trans-cellular tissue system. Together with adherens junctions and tight junctions, desmosomes enable integration and stratification of epithelia, whilst ensuring that barrier function is maintained at all times and also that damaged tissue cells can be replaced. Loss of desmosome function is linked to severe diseases, particularly in mechanically challenged tissues such as skin and cardiac muscle (reviewed by (Chidgey and Dawson, 2007; Delmar and McKenna, 2010; Dusek and Attardi, 2011; Ishida-Yamamoto and Igawa, 2014; Maruthappu et al., 2019; Spindler et al., 2018; Thomason et al., 2012).

Within the desmosomal macromolecular complex, the extra-cellular domains of the desmosomal cadherins (DCs), desmocollins (Dsc) and desmogleins (Dsg), bind tightly to their counterparts on the adjacent cells, while their intracellular domains connect via plakoglobin (PG) and plakophilins (Pkps) to intermediate filament–binding dimers of desmoplakin (DP) (Garrod and Chidgey, 2008; Green et al., 2019). Desmosomes of intact homeostatic tissues exist in a state of hyper-adhesion, rendering them particularly stress-resistant (Wallis et al., 2000; Garrod et al., 2005). Whilst newly-formed desmosomes need extracellular calcium for adhesion through their DCs, mature hyper-adhesive desmosomes remain robustly attached to each other even after experimental removal of extracellular calcium by experimental chelation of calcium ions (Ca^2+^).

Hyper-adhesion is developmentally regulated in the mouse embryo and is acquired with time in confluent culture of several cell lines (Kimura et al., 2007; Kimura et al., 2012; Wallis et al., 2000). It has been suggested that hyperadhesion is associated with an ordered arrangement of the extracellular domains of the desmosomal cadherins, indicated by the presence of an electron-dense midline present halfway between the desmosomal cell membranes, and that this may result in Ca^2+^ being locked into the structure (Garrod et al., 2005). It has been reported that this ordered arrangement of extracellular domains of genetically modified Dsg3 is lost upon removal of extracellular Ca^2+^, both from pharmacologically-induced hyper-adhesive and calcium-dependent desmosomes, suggesting the ordering of DCs is not the only factor involved in desmosomal adhesion (Bartle et al., 2020; Bartle et al., 2017).

Whilst tissue integrity depends upon the maintenance of stable cell-cell junctions, cells must retain the ability to free themselves from their neighbours in order to move or to get rid of damaged cells. During embryonic development, in wound healing and tissue regeneration, cell adhesion must be temporarily relaxed or lost for cells to undertake tissue remodelling. This is seen in epithelial-to-mesenchymal transition (EMT), the conversion of compact epithelial sheets to solitary or small groups of motile cells, which occurs during development and in metastatic progression of some cancers, and is often triggered by growth factor signalling (Nieto et al., 2016).

Despite the importance of cell adhesion in health and disease, little is known about how desmosomal adhesion is lost in vivo. A limited amount of evidence suggests that loss of desmosome-dependent cell adhesion may occur through internalisation of whole desmosomes by the cells involved (Allen and Potten, 1975; Garrod et al., 2005). This has not been studied in detail because no tissue culture model of the process exists. Studies on the internalisation of half desmosomes, induced by (non-physiological) Ca^2+^ chelation, have shown that this process is actin dependent, that the internalised half-desmosomes remain intact until degraded by a combination of lysosomal and proteasomal activity, and that desmosomal proteins were not recycled (McHarg et al., 2014). Whether similar mechanisms apply to the internalisation of whole desmosomes under physiological conditions remains to be determined.

Desmosomes are undoubtedly highly stable structures, stability being key to their function, and it is becoming clear that the protein exchange rates within desmosomes mirror their stability as has been shown for Dsg2, Dsg3, Dsc2, DP and PG (Bartle et al., 2020; Foote et al., 2013; Gloushankova et al., 2003; Lowndes et al., 2014; Moch et al., 2019; Vielmuth et al., 2018; Windoffer et al., 2002). Nevertheless they are also dynamic, with a life cycle of four phases: de novo assembly from their component molecules, a weakly-adhesive calcium-dependent phase compatible with cell migration, a strongly adhesive calcium-independent phase in tissue homeostasis, and desmosome removal or breakdown as epithelia become activated by relevant signalling.

We investigated the complex relationships between desmosomal proteins during the last three stages of this life cycle using time-lapse fluorescence microscopy including fluorescence recovery after photobleaching (FRAP) and fluorescence loss and localisation after photoactivation (FLAP). A culture model for desmosome internalisation was established to mimic the in vivo situation more closely, and we observed that one of the desmosomal proteins, plakophilin 2a (Pkp2a), behaves very differently from the others. In calcium-dependent desmosomes the molecular mobility of Dsc2a, Dsg2 and PG is very low and is further reduced as the cells become hyper-adhesive, forming a stable desmosomal core. DP is mobile initially but stabilises with the core with the onset of hyper-adhesion. In contrast, Pkp2a shows a unique and persistently dynamic exchange between its desmosomal and a cytoplasmic pools. Similar relative protein dynamics persist when whole desmosomes are internalised during HGF-induced scattering of Madin-Darby canine kidney (MDCK) cells. These results suggest a uniquely dynamic role for Pkp in desmosomes.

## Results

### Calcium-dependent desmosomes exhibit differential protein dynamics

To shed light on the behaviour of the individual desmosomal components, we first analysed their dynamic properties using FRAP. The relative mobility of the proteins was assessed by determining their half-time of fluorescence recovery (t_1/2_) and mobile fraction (F_m_) (Carisey et al., 2011). Dsc2a, Dsg2, PG, DP and Pkp2a tagged with either neonGreen or eGFP fluorophores were stably expressed in the simple epithelial cell line MDCK type II (Fig. 1 A), a well-established model for the study of cell-cell junctions (Dukes et al., 2011). The cells were plated at confluent density and cultured for 24h prior to FRAP, at which time the majority of the desmosomes showed calcium-dependent adhesion (Fig. S1 A, B) (Wallis et al., 2000).

**Fig. 1.**
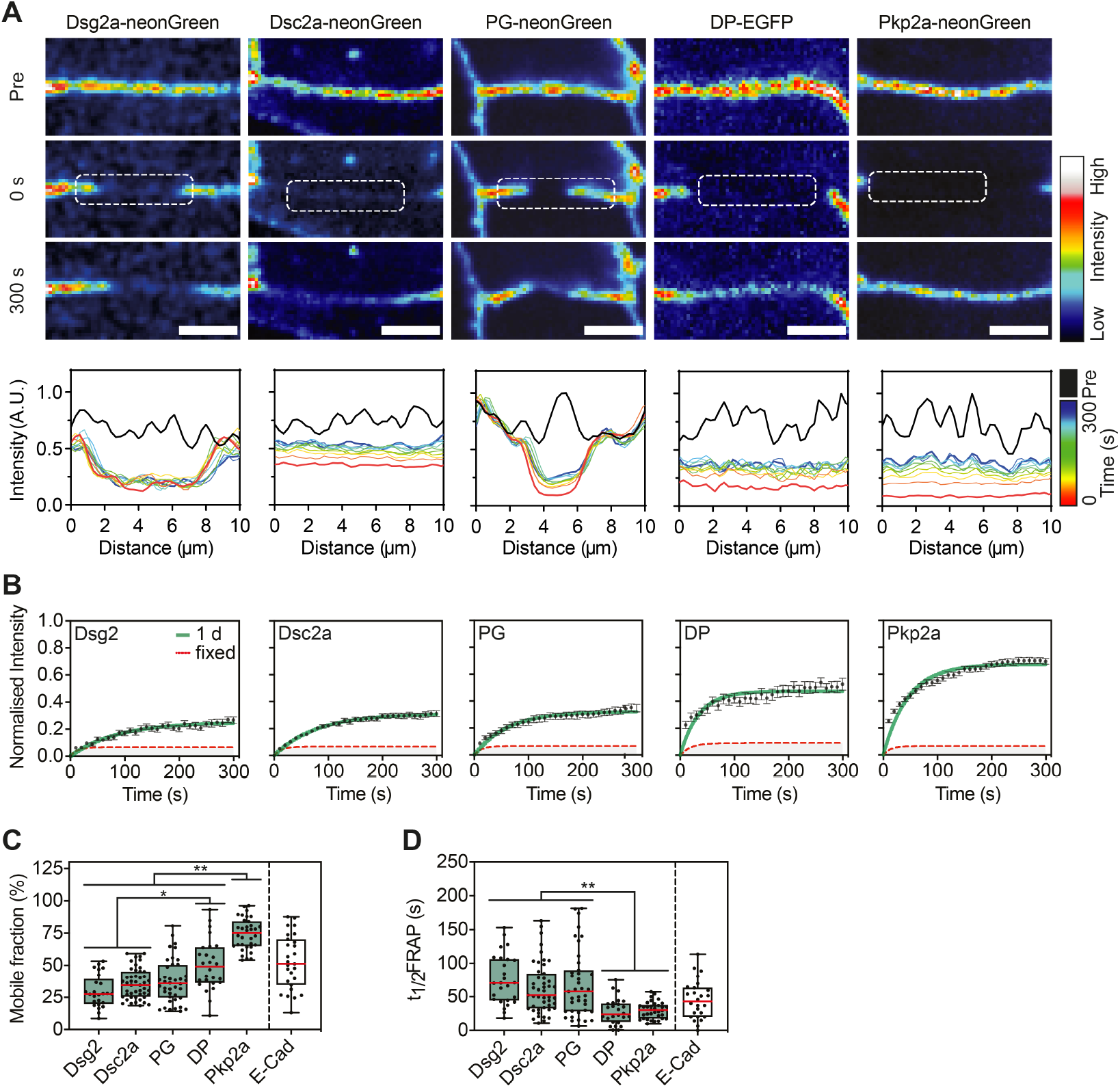
Calcium-dependent desmosomes exhibit differential protein dynamics. **A**. Representative time-lapse images showing FRAP in MDCK cells transfected with neonGreen-labelled Desmoglein 2 (Dsg2), Desmocollin 2a (Dsc2a), Plakoglobin (PG) and Plakophilin2a (Pkp2a), and EGFP tagged Desmoplakin (DP). Images are presented in a colour intensity scale. Graphs show the line profile plots of the bleached ROIs (dashed box at 0 s) over time in a colour intensity scale. Scale bar: 5 µm. Note the recovery of Pkp2a in similar spots as pre-bleaching. **B**. Graphs show the mean fluorescence recovery curves for all recorded desmosomes (n=24-47, N=3); red line showing the recovery of PFA-fixed samples; error bars are s.e.m. **C**. Mean mobile fraction and **D**. t_1/2_ FRAP values for the indicated desmosomal proteins in MDCK cells (scatter plot). n=24-4, N=3. The box represents the 25-75th percentiles, and the median is indicated in red. The whiskers show the range of values. **P 0.01; *P 0.05 (Kruskal-Wallis with Dunn’s multiple comparisons test).

A wide distribution in the (F_m_) of the desmosome proteins was observed (Fig. 1 B). The transmembrane proteins Dsg2 (F_m_=29.8%) and Dsc2a (F_m_=35.7%) showed the lowest mobilities, closely followed by the cytoplasmic plaque protein PG (F_m_=37.7%). DP was slightly more mobile (F_m_=50.2%), while Pkp2a showed the highest mobility (F_m_=74.6%) (Fig.1C). Analysis of the recovery times showed similar results, with slower recovery times of DCs (Dsg2 t_1/2_=75.7s, Dsc2a t_1/2_=62.9s) and PG (t_1/2_=60.1s) in comparison to significantly faster ones for DP (t_1/2_=27.6s) and Pkp2a (t_1/2_=30.1s) (Fig. 1 D). To put the high stability of the DCs into perspective against actin-associated cell-cell junction proteins, we measured the mobility of E-cadherin, the adhesion receptor of adherens junctions. Mobility of E-cadherin tagged with mEmerald (E-Cad) was much higher (F_m_=52.8%) than the mobility of the DCs, demonstrating that desmosomal receptors are more stable at the cell-cell interface than E-cadherin in adherens junctions (Fig. 1 C).

The very low mobile fraction of Dsg2, Dsc2a and PG prompted us to evaluate the contribution of the reversible photobleaching properties of the fluorophores, which can be up to 20% of the fluorescence intensity prior to the bleaching event (Sinnecker et al., 2005). Performing FRAP on paraformaldehyde fixed cells we recorded a F_m_ of 6.8% for neonGreen and a F_m_ of 10.2% for eGFP after 5 minutes of recordings, which readings are related to the inherent properties of the fluorescent tags (red dotted line Fig. 1 B). These recovery rates suggested an even lower mobility of the desmosomal proteins. However, these values cannot directly be subtracted from the F_m_ measured in the live cell recordings as the fixation process may lead to conformational changes in the fluorophores that may result in varied lifetimes and properties (Becker, 2012).

To bypass the potential issue of reversible photobleaching and for further analysis, we analysed FLAP (Fig. 2 A). This allowed us to calculate the fluorescence decay within the activated region of interest (ROI) as well as to track the activated fluorescent proteins within their environment. Using photoactivatable-GFP (PaGFP) tagged Dsc2a, with mScarlet-Pkp2a used as a desmosome marker, and PG-PaGFP with Dsc2a-mScarlet, FLAP showed similar protein turnover readings to those found by FRAP. Both techniques showed a low mobile fraction while Pkp2a-PaGFP had a significantly higher turnover as seen by FRAP (Fig. 1 C, 2 B). We conclude that in calcium-dependent desmosomes, DCs and PG form a stable desmosomal core, whereas Pkp2a has a dramatically higher mobility that suggests a more transient association of this protein with desmosomes. In calcium-dependent desmosomes the mobility of DP is intermediate between that of the core and the highly mobile Pkp2a.

**Fig. 2.**
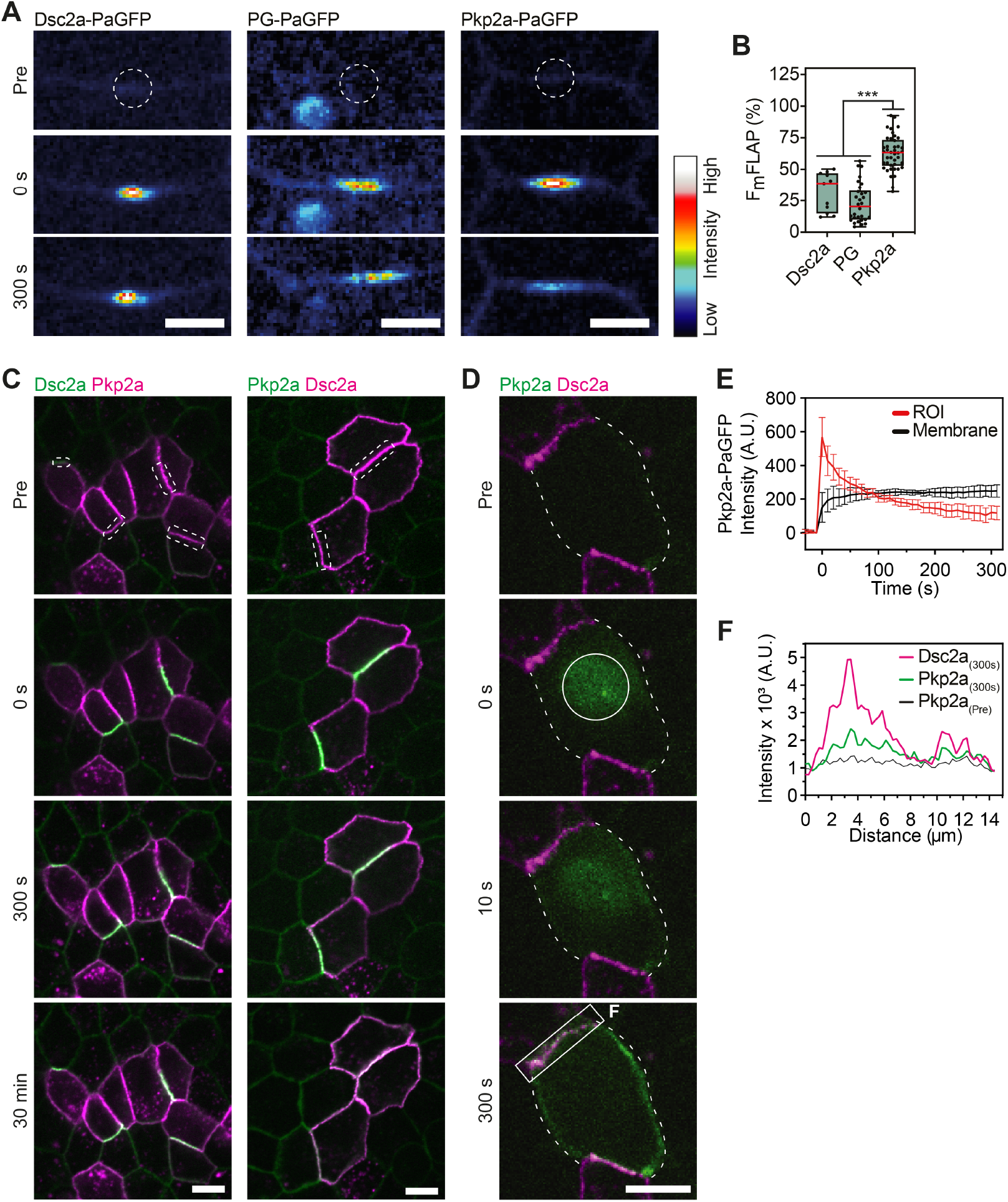
Cytoplasmic pool of plakophilin 2a enables rapid exchange at desmosomes. **A**. Representative time-lapse images showing FLAP in MDCK cells transfected with PaGFP tagged Desmocollin 2a (Dsc2a), Plakoglobin (PG) and Plakophilin2a (Pkp2a). Images are presented in a colour intensity scale. Scale bar: 5 µm.**B**. Mean FLAP mobile fraction of Dsc2a, PG and Pkp2a. The box represents the 25-75th percentiles, and the median is indicated in red. The whiskers show the range of values. ***P 0.001 (student’s t-test).**C**. Time-lapse images showing photoactivation of Dsc2a-PaGFP in comparison to PaGFP-Pkp2a at desmosomes over a 30 min period. Scale bar: 10 µm.**D**. Cytoplasmic photoactivation of PaGFP-Pkp2a indicated by the 10 µm circular ROI at 0s. The outline of the cell is indicated by the dotted lines. Scale bar: 10 µm. **E**. Fluorescence intensity of PaGFP-Pkp2a over time showing the rapid decrease in photoactivated cytoplasmic ROIs concomitant signal increase at the membrane (N=3). **F**. Line profile plots showing overlapping peaks of fluorescence signal of PaGFP-Pkp2a and Dsc2a-mScarlet after 5 min indicated by the box in **D**.

### Plakophilin 2a shows rapid exchange between desmosomes and a cytoplasmic pool

The mobility of desmosomal proteins could be due to either a local exchange of proteins at the plasma membrane or due to an exchange with intracellular pools, or both. In order to determine the mechanism of Pkp2a exchange at desmosomes, we performed further FLAP experiments. FLAP recordings of PaGFP-Dsc2a activated at the plasma membrane were taken over 30 mins, which confirmed the extraordinarily stable localisation of Dsc2a (Fig. 2 C). In contrast, photoactivated Pkp2a-PaGFP at the same settings distributed rapidly along the entire membrane (Fig. 2 C). To assay the flux between the cytoplasmic pool of Pkp2a and the desmosome-associated Pkp2a, we performed FLAP by targeting the 405nm laser light to a 10 µm diameter circular region of the cytoplasm for localized photoactivation and tracking of PaGFP-Pkp2a out from this region (Fig. 2 D). PaGFP-Pkp2a diffused within seconds throughout the entire cytoplasm and was found to become rapidly enriched at the membrane (Fig. 2 D, E). Line profile analysis revealed overlapping fluorescence intensity peaks of PaGFP-Pkp2a and Dsc2a-mScarlet, indicating the recruitment of Pkp2a from the cytoplasm to desmosomes (Fig. 2 F). Similar FLAP attempts with the other desmosomal components of Dsc2a and PG failed because of their extremely low presence in the cytoplasm (Fig. S1 C).

Taken together, the results revealed a previously unknown property of Pkp2a, i.e. that it circulates continuously between the desmosomal plaque and the cytoplasm, in stark contrast to the more stable association of the other studied components with the desmosomes.

### Hyper-adhesive desmosomes are more stable yet retain high Pkp2a turnover

We next asked whether the dynamics of the major desmosomal proteins changed as desmosomes matured from calcium-dependent to calcium-independent hyper-adhesion, with two major questions in mind: (a) do the major proteins acquire greater stability, and (b) does Pkp2a remain more dynamic than the other proteins?

We first established in all the cell lines we tested that most desmosomes had become hyper-adhesive after 3 days of confluent culture (Fig. S1 A, B). To investigate whether the adhesive state of desmosomes affects their protein dynamics, we performed FRAP in 3-day confluent MDCK cells (Fig. 3 A, B) and compared them with those obtained in the 1-day cultures (Fig. 1). All the tested desmosomal proteins became significantly less mobile as desmosomes matured to hyperadhesion (Fig. 3 B, Table 1). In contrast, the mobility of E-cadherin was not significantly reduced over the same time period. Dsg2 showed the lowest mobile fraction of 13.6% followed by Dsc2a and PG (both <25%). Taking into account reversible photobleaching of up to 10% showed that these three proteins remained almost static in cell-cell junctions. The low mobile fraction rendered it impossible to determine a reliable t_1/2_ for the proteins due to potential interference from rapidly reversible photobleaching seen in fixed cells (neon-Green t_1/2_=11.3s, EGFP t_1/2_=29.8s). A significant decrease of mobility was also observed for DP and Pkp2a (Fig. 3 C and Table 1). DP showed the greatest decrease in its mobile fraction from 50.2% to 30.0% concomitant with a reduction in the t_1/2_. Although Pkp2a also showed decreased mobility under hyper-adhesive conditions, it retained a high mobility of 58.8%. This was at least twice the level of mobility of the other desmosomal proteins and was particularly striking when line profiles of the bleached regions were compared (Fig. 3 A). These results were confirmed by FLAP (Fig. 3 D, S1 D). The mobility of both Dsc2a and PG tagged with PaGFP was significantly reduced after 3 days of confluent culture but Pkp2a retained a considerably higher turnover. These results demonstrate a substantial decrease in mobility of the desmosomal components accompanying maturation to hyper-adhesion. Furthermore, they support the idea of a desmosomal core in which DP joins the core formed by DCs and PG at earlier stages. Pkp2a, however, has a novel and surprisingly dynamic role.

**Fig. 3.**
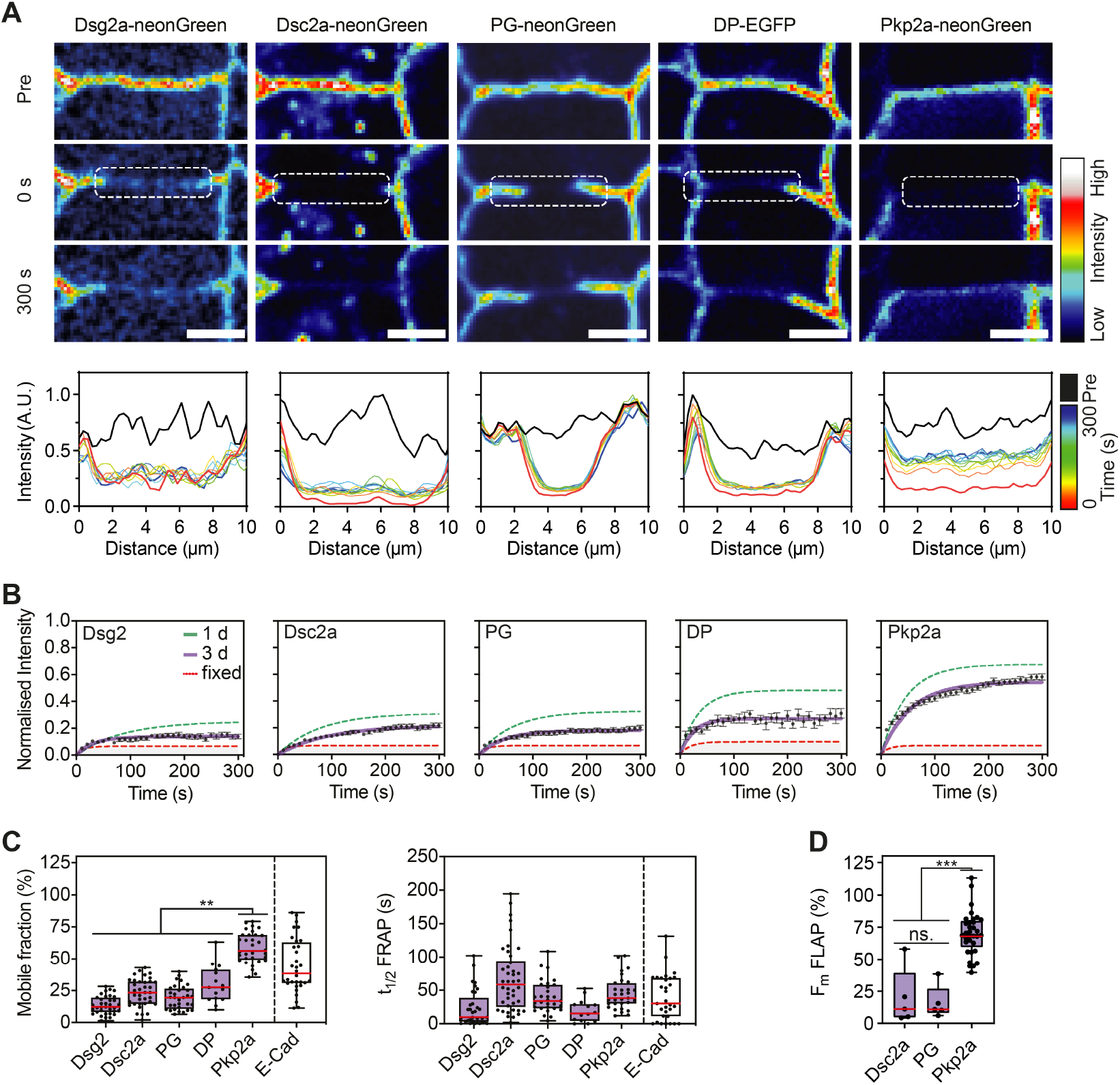
Protein dynamics are reduced but Pkp2a remains significantly more mobile in hyper-adhesive desmosomes. **A**. Representative time-lapse images showing FRAP in MDCK cells transfected with neonGreen-labelled Desmoglein 2 (Dsg2), Desmocollin 2a (Dsc2a), Plakoglobin (PG) and Plakophilin2a (Pkp2a), and EGFP tagged Desmoplakin (DP). The cells were cultured confluent for 3 d leading to the acquisition of hyper-adhesion in the majority of desmosomes. Images are presented in a colour intensity scale. Representative time-lapse images showing FRAP of MDCK cultured confluent for 3 days. Graphs show the line profile plots of the ROIs over time in a colour intensity scale. Scale bar: 5 µm. **B**. Graphs show the mean fluorescence recovery curves for all recorded desmosomes in 3 d monocultures (n=15-42); dashed green line indicates mean recovery curve in 1 d monocultures; error bars are s.e.m. **C**. FRAP mean mobile fraction and t_1/2_ FRAP values for the indicated desmosomal proteins in 3 d monolayers (scatter dots). (n=15-42, N=3). **D**. FLAP mean mobile fractions of Dsc2a, PG and Pkp2a comparing 1d or 3d cultured monolayers. The box represents the 25-75th percentiles, and the median is indicated in red. The whiskers show the range of values. ns. no significant difference; *P<0.05; **P<0.01; ***P<0.001 (Kruskal-Wallis with Dunn’s multiple comparisons test).

### Intact desmosomes are internalised during cell scattering

The existence of a stable fraction of core desmosomal components, even where the binding of the DCs was calcium-dependent, led us to question whether this would be maintained during desmosome down-regulation, which is required for morphogenesis, wound healing, tissue remodelling and EMT. Alternatively, would desmosomal components become highly mobile leading to desmosome dissolution?

In order to create a culture model of desmosome down-regulation that did not require Ca^2+^ chelation, we used HGF-induced scattering of MDCK cells, which leads to phenotypic changes resembling partial EMT, including increased cell motility and loss of cell-cell adhesion (Balkovetz, 1998; Ridley et al., 1995). Desmosomes were tracked using realtime imaging of mixed populations of MDCK cells expressing either Dsc2a-YFP or Dsg2-mCherry. Thus, where two differentially-labelled cells were in contact, desmosomes were labelled with Dsc2a-YFP (green) on one side and Dsg-mCherry (magenta) on the other (Fig. 4 A, C). To test the validity of this system we first treated such mixed populations with Ca^2+^ chelation which has been well documented to induce splitting of desmosomal adhesion and internalisation of half desmosomes. Imaging showed the extracellular separation of the desmosomes into two half-desmosomes which were subsequently internalised by their own cell as indicated by the entry of solely magenta or solely green particles into the cells (Fig. 4 A, C). These observations were consistent with the results of previous Ca^2+^ chelation experiments in which half-desmosomes were seen to be internalised (Mattey and Garrod, 1986; McHarg et al., 2014).

**Fig. 4.**
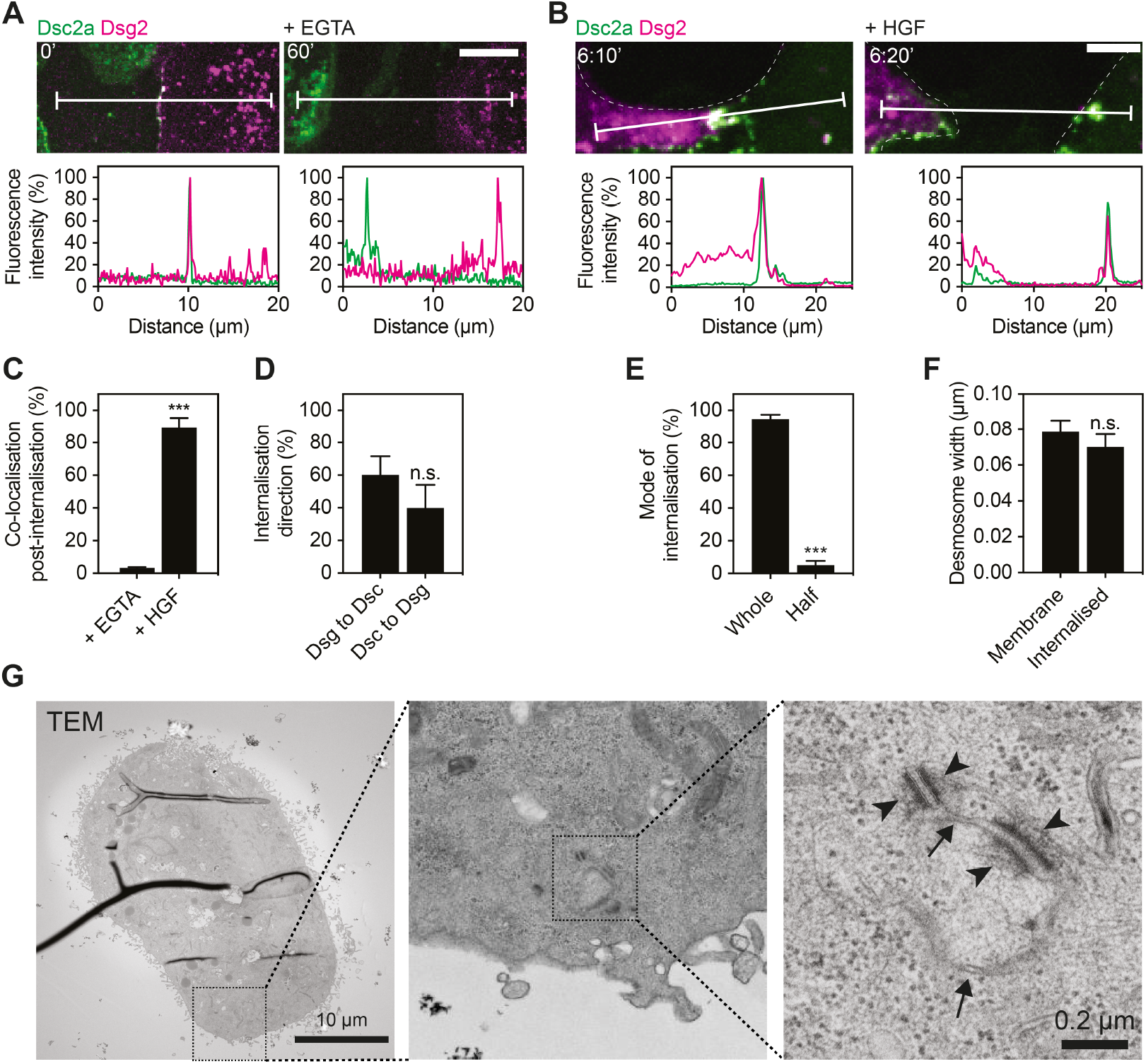
Hepatocyte growth factor-induced cell scattering leads to whole desmosome internalisation. **A**. Mixed populations of MDCK cells expressing either Dsc2a-YFP (green) or Dsg2-mCherry (magenta) were cultured in SM for 1 d. Desmosomes are indicated by the co-localisation of the fluorescence signals. Separation of the cells following Ca^2+^ chelation (+EGTA) for 60 minutes led to the separate internalisation of Dsc2-YFP and Dsg2-mCherry.**B** Whereas cells treated with 40 ng/ml hepatocyte growth factor (HGF) induced scattering leading to the joined internalisation of both Dsc2a-YFP and Dsg2-mCherry by one or the other of the adjacent cells. Scale bar: 5 µm. Fluorescence line profiles depict the fluorescence intensity along the white line in the images.**C** Quantification of co-localisation of Dsc2a-YFP and Dsg2-mCherry after internalisation induced by Ca^2+^ chelation or HGF treatment.**D** Quantification of internalisation direction either Dsg2-mCherry into the Dsc2a-YFP expressing cell or vice versa following HGF treatment.**E** Analysis of the mode of desmosome internalisation, either whole or half, revealed by transmission electron microscopy (TEM) following HGF treatment.**F** Graph showing the plaque to plaque width of the desmosomes measured on EM images remains mostly unchanged after internalisation. (n = 10 desmosomes per group) **G** Representative TEM images of internalised double membrane structures (arrows) with intact whole desmosomes with attached intermediate filament fragments on each side of the doublet (arrowheads). ***P<0.001; n.s. no significant difference (student’s t-test).

In order to monitor the down-regulation of desmosomes in the presence of a physiological calcium concentration, such mixed cell populations were grown on fibronectin-coated glass and induced to scatter by treatment with HGF following serum starvation. Under these conditions, cells growing in small islets started spreading and single cells detached from their neighbours (Fig. S2 A). High magnification fluorescence imaging at 5-minute intervals revealed that desmosomes remained stable at the sites where cells maintained contact. Surprisingly, during the final stages of cell separation, whole desmosomes were internalised intact by one or other of the pair of adjacent cells (Fig. 4 B, C), with the entire desmosomal structure becoming engulfed, including cytoplasmic fragments from both cells (i.e. fluorescence signals of Dsc2a-YFP and Dsg2-mCherry). There was no preference for whether the Dsc2a-YFP or the Dsg2-mCherry expressing cell internalised the whole desmosomes (Fig. 4 D). Quantification of the co-localisation of fluorescent particles after internalisation revealed that concomitant internalisation of the Dsg and Dsc complex from opposing cells was the dominant mode of internalisation following HGF treatment (Fig. 4 C). These results implied that HGF-induced scattering led to the internalisation of whole desmosomes, rather than separation of the desmosomal receptors from neighbouring cells or the complete dissolution of desmosomes. To confirm this hypothesis, we performed electron microscopy (EM). We used correlative light and electron microscopy (CLEM) to identify cells with internalised desmosomes (Fig. S2 B). Subsequent ultrastructural analysis of the EM micrographs showed that the internalised, double-labelled structures were indeed morphologically intact desmosomes with attached keratin filaments and remains of the plasma membrane (Fig. 4 G). Comparing the width of desmosome plaques as a readout for their intactness showed only a slight reduction from those of intact desmosomes at the membrane (Fig. 4 F). The great majority (>90%) of desmosomes internalised following HGF treatment were internalised whole, with a small number of structures identified as half-desmosomes (Fig. 4 E).

We concluded that desmosomes were predominately mechanically removed without undergoing any form of dissolution. Rather, they appear to be torn away from one cell and internalised by its neighbour.

### Internalised desmosomes include all major desmosome proteins plus keratin filament fragments

To determine which of the multiple components remained associated with internalised desmosomes after cell separation, we examined their composition using fluorescence super-resolution microscopy following immunostaining for major desmosomal proteins. Structured illumination microscopy (SIM) and co-localisation analysis revealed that the majority of internalised desmosomes maintained their protein composition following internalisation (Fig. 5 A, B). The high resolution of SIM clearly revealed the two distinct patches of DP in the intact desmosomes (Fig. 5 A, lower left panel). This morphology, referred to as “railroad tracks” (Stahley et al., 2016), was also observed post internalisation (Fig. 5 A). Analysis based on images taken by wide field microscopy showed internalised Dsc2a-YFP co-localised with DP (81% of all internalisation events), PG (91%), Pkp2a (72%) and keratin 8 (KRT8, 88%) (Fig. 5 B). These results support the interpretation that the internalised desmosomes were intact, both at the morphological level and in terms of protein composition. Electron micrographs showed intermediate filaments attached on both sides (plaques) of the internalised desmosomes (Fig. 4 G). Also SIM analysis suggested that internalised desmosomes were still attached to keratin filaments after internalisation (Fig. 5 A, lower right panel). Some of the high resolution images suggested that keratin filaments were severed at one side of the desmosome originating from the neighbouring cell (Fig. 5 C). To examine this possibility further, we carried out live-cell imaging on cells stably expressing Dsc2a-YFP which were transiently co-transfected with mCherry-KRT18. This resulted in all cells possessing YFP-labelled desmosomes but only a subset showing mCherry-labelled keratin filaments. Where pairs of mCherry-KRT18-expressing cells with K18-untransfected cells were tracked through HGF-induced scattering, co-internalisation of mCherry-KRT18 with Dsc2a-YFP into the non-transfected cell was observed (Fig. 5 D). These results are consistent with the view that the force involved in cell separation results in rupture or severing of the keratin filaments in one of the opposing cells, rather than pulling the desmosome plaques apart from each other. This reinforces the picture of desmosomes as extremely stable cell-cell attachments.

**Fig. 5.**
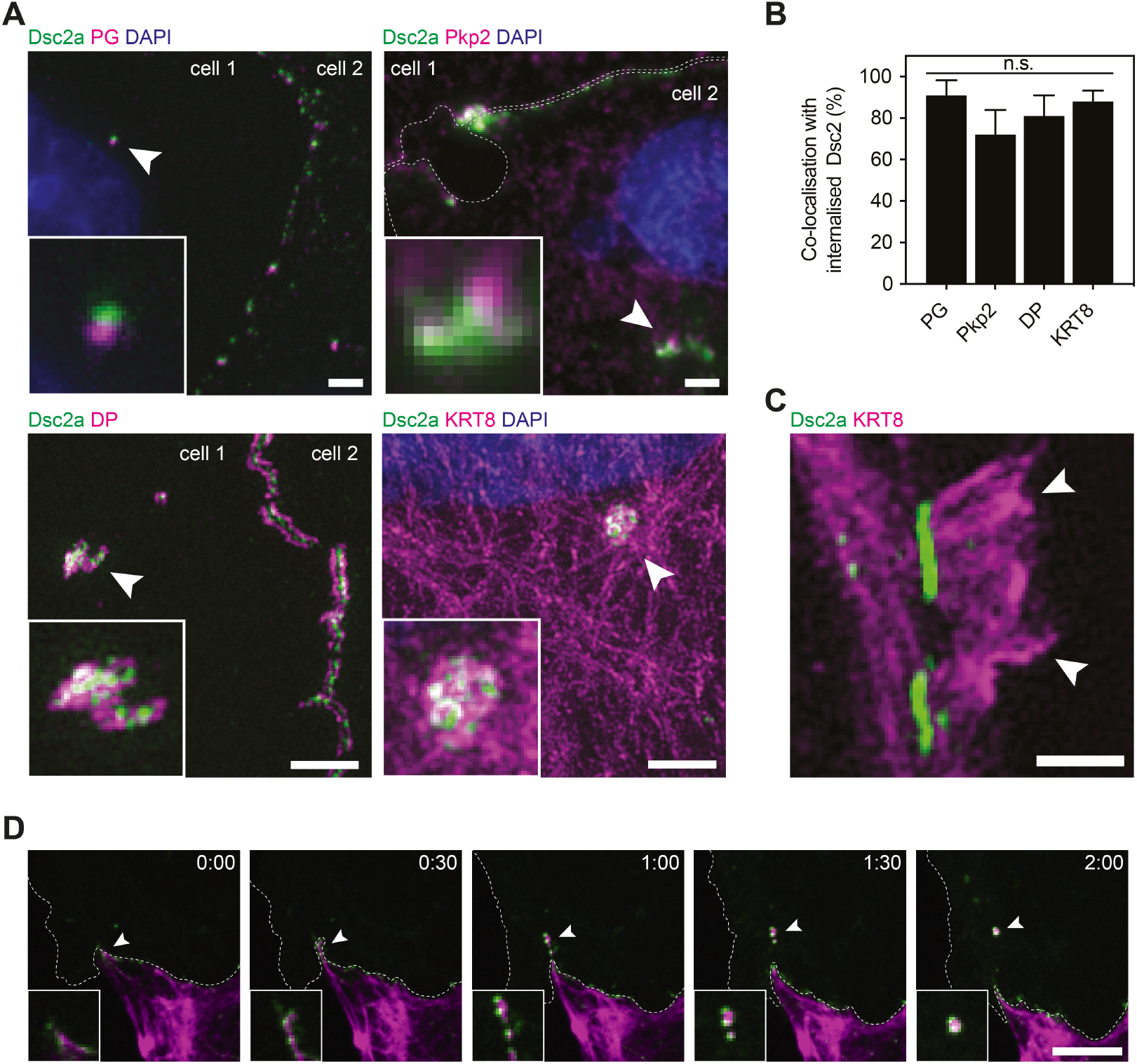
Internalised whole desmosomes are composed of all major desmosomal and keratin fragments. **A**. MDCK cells expressing Desmocollin 2a tagged with YFP (Dsc2a) were induced to scatter by HGF treatment. Cells were immunofluorescence stained for desmoplakin (DP), plakophilin 2 (Pkp2), plakoglobin (PG) and keratin filaments (KRT8). Representative structured illumination microscopy (SIM) images of internalised desmosomes. Scale bar: 5 µm.**B** Quantification of the co-localisation of internalised Dsc2a with the respective protein.**C** Representative SIM image showing fragments of keratin filaments attached to internalised desmosomes. Scale bar: 1 µm.**D** Time-lapse images of MDCK cells expressing either Dsc2a-YFP (green) KRT8-mCherry (magenta) treated with HGF demonstrate the internalisation of keratin fragments of the adjacent cell. Scale bar: 5 µm.

### Desmosome protein mobilities remain consistent after internalisation

Since desmosomes remained structurally intact following internalisation, we next asked whether internalisation changed the stability or the dynamic behaviour of the individual desmosomal components. We therefore analysed FLAP on the desmosomal core component Dsc2a and the putative signalling component Pkp2a in internalised desmosomes after 4h of HGF treatment. Both Dsc2a as part of the desmosomal core and Pkp2a showed a comparable FLAP profile following HGF treatment to the PBS control (Fig. 6). The results indicate that the functional structure of desmosomes, encompassing both the stable core and the transiently interacting Pkp2a, remained consistent after internalisation. It remains to be determined whether these internalised structures therefore perform any further function within the cells.

**Fig. 6.**
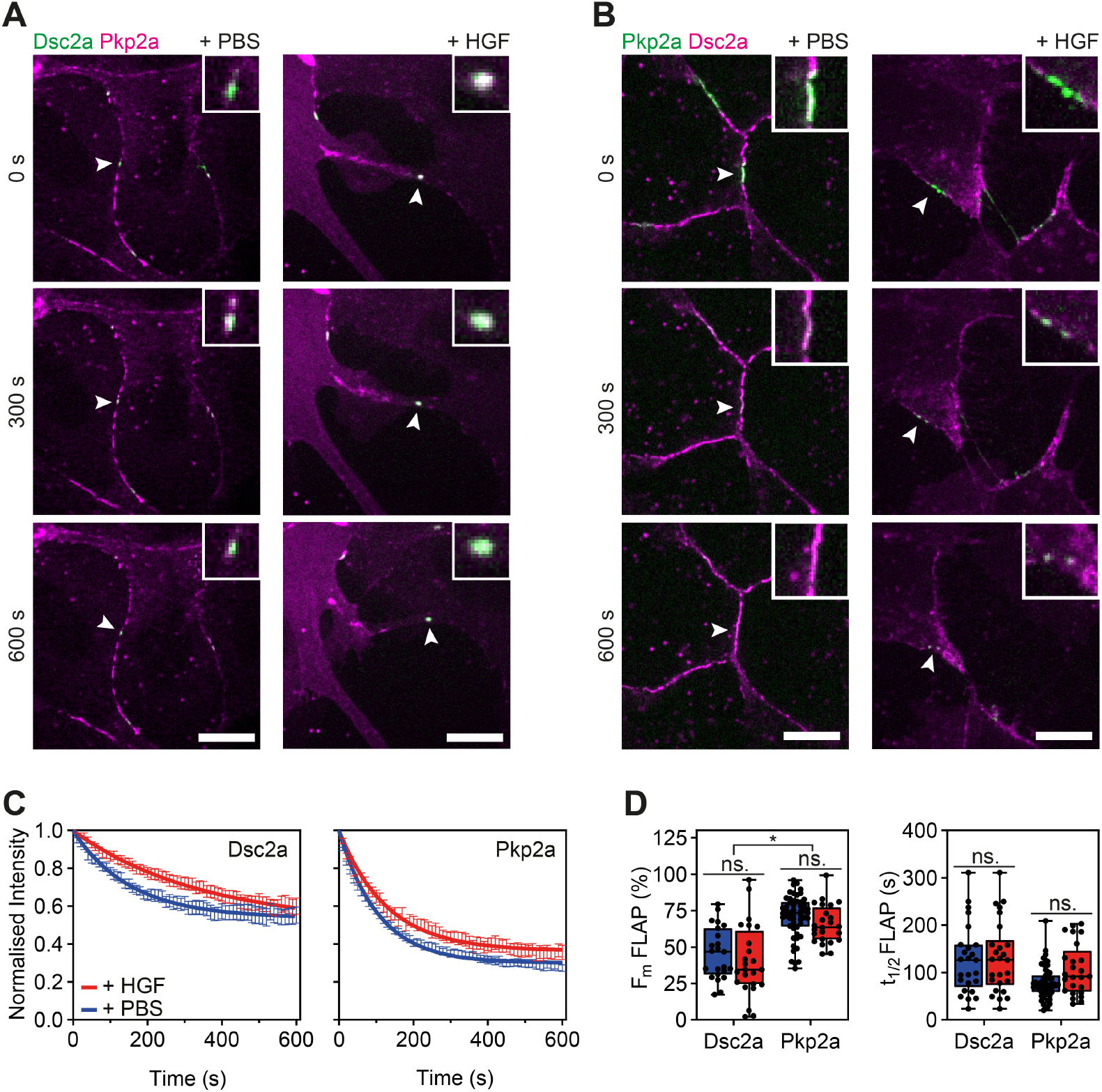
Mobility of desmosomal proteins remains unchanged following internalisation. **A** Representative time-lapse images showing FLAP in MDCK cells transfected with Dsc2a-PaGFP and mScarlet-Pkp2a. Cells were cultured were treated for 4 h with 40 ng/ml hepatocyte growth factor (HGF) or same v/v PBS.**B** In comparison to **A** MDCK cells were transfected with PaGFP-Pkp2a and Dsc2a-mScarlet. Scale bar: 10 µm.**C** Graphs show the mean fluorescence decay curves for all recorded desmosomes (n=23-53, N=3); error bars are s.e.m.**D** FLAP mean mobile fractions and t_1/2_ of Dsc2a and Pkp2a comparing PBS control and HGF treated cells. The box represents the 25-75th percentiles, and the median is indicated in red. The whiskers show the range of values. ns. no significant difference; *P<0.05 (Kruskal-Wallis with Dunn’s multiple comparisons test).

## Discussion

Our major finding is that desmosomes consist of two contrasting protein fractions or modules. These are: (1) a strikingly stable core module, composed of the DCs and PG, at the calcium-dependent stage, and (2) a high mobility module, represented by Pkp2a, which exhibits a transient interaction with the desmosome core and a fast turnover between its desmosome-associated pool and a cytoplasmic pool. Both modules become less dynamic as desmosomes mature from calcium-dependence to hyper-adhesion. While DP transits to behave more like a stable core molecule at this stage, Pkp2a retains high molecular mobility. This modular composition of desmosomes is maintained even during growth factor-induced epithelial cell separation when whole double-plaque-bearing desmosomes are internalised by the scattering cells (Fig. 7). We further demonstrate that these internalised desmosomes are fully intact and even retain attached, apparently torn keratin filaments from the formerly adjacent cell. This suggests that force is involved during the loss of cell-cell adhesion and cell separation. It appears that the cells are literally torn apart by the force generated by their scattering movement.

**Fig. 7.**
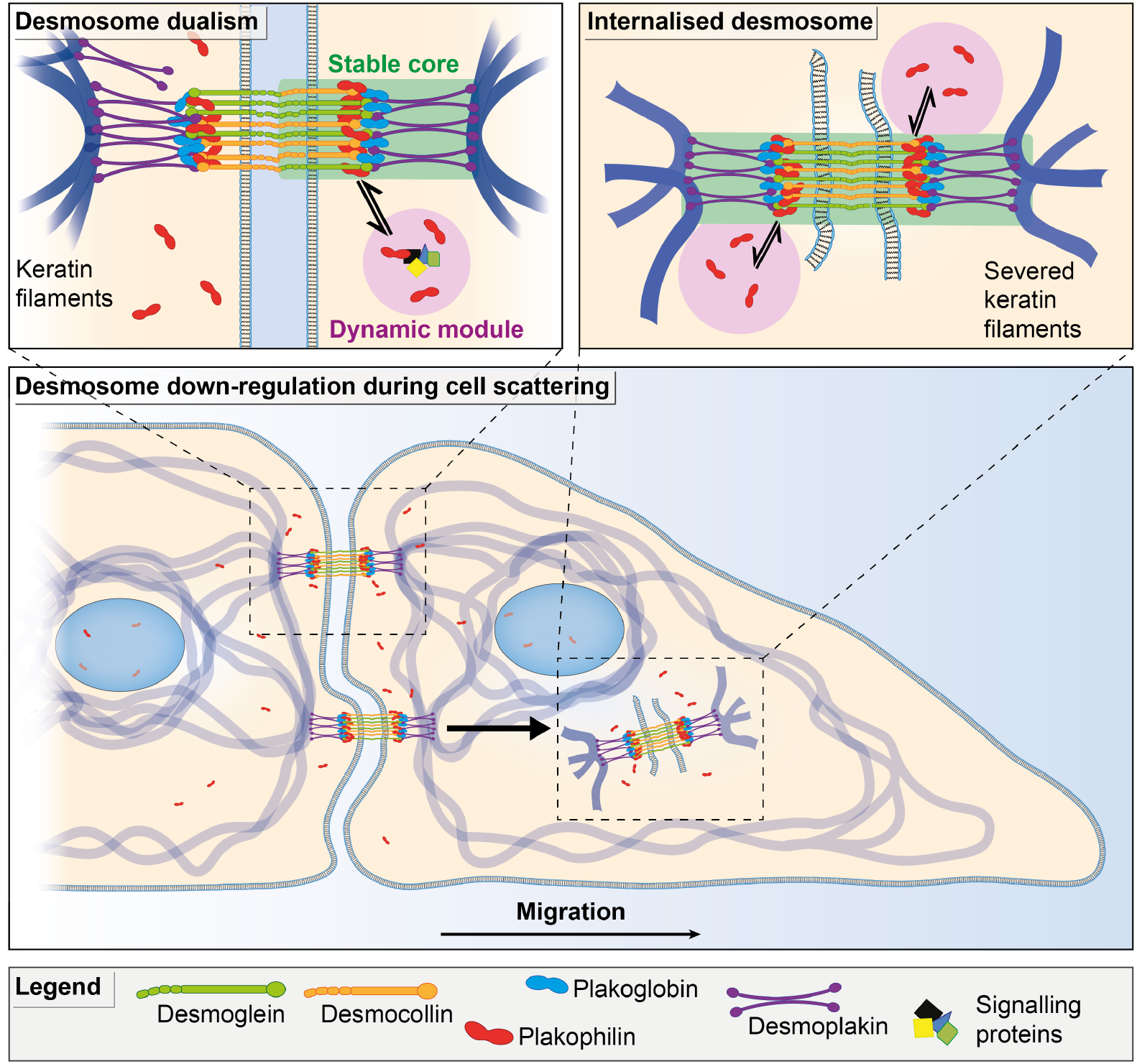
Model of desmosome dualism and down-regulation during growth factor-induced cell separation. Desmosomes consist of two contrasting protein fraction, a strikingly stable core, composed of the DCs and PG, at the calcium-dependent stage, and a high mobility module, represented by Pkp2a. As desmosomes mature from calcium-dependence to calcium-independent hyper-adhesion, core stability increases, but Pkp2a remains highly mobile. Desmoplakin mobile initially, stabilises with hyper-adhesion. Desmosome down-regulation during growth factor-induced cell scattering proceeds by internalisation of whole desmosomes, which still retain a stable core and highly mobile Pkp2a.

FRAP and FLAP allow detailed analysis of the integration of proteins in subcellular compartments (Ishikawa-Ankerhold et al., 2012). Our study has revealed a diverse profile of protein dynamics of components within desmosomes that are engaged in cell-cell adhesion, the most striking observation being that Pkp2a is highly dynamic throughout desmosome maturation and downregulation, while the DC adhesion receptors are exceptionally stable. This finding is even more surprising because ultrastructural studies have shown that Pkp is located in the outermost region of the outer dense desmosomal plaque (Al-Amoudi et al., 2007; North et al., 1999). Our results suggest that molecular exchange can take place readily within this apparently dense and well organised structure. For integrin mediated cell-matrix adhesions (focal adhesions), it was previously shown that structural proteins that directly link adhesion receptors to the contractile actin cytoskeleton are less mobile than proteins controlling fast signalling processes (e.g. GTPase activity) (Stutchbury et al., 2017). By analogy, the interpretation of our data in this study may lead to a model whereby the DCs, PG and DP form the main stable axis to the intermediate filament (IF) network and that Pkp can act as the major signalling component distributing information derived from desmosomes to other compartments of the cell. Such model would be in line with the multiple ways Pkps are involved in a variety of signalling events (Hatzfeld, 2007; Hatzfeld et al., 2014). Two examples that may particularly benefit from fast translocation of Pkp2 to different compartments are its reported roles in regulating RhoA localisation thus mediating signals that also affect the actin cytoskeleton, and in transcriptional regulation in the nucleus (Cerrone et al., 2017; Godsel et al., 2010; Mertens et al., 1996; Sobolik-Delmaire et al., 2010). There is increasing evidence that this occurs in collaboration with components of adherens junctions and the fast cycling Pkp2, potentially through competitive binding with E-cadherin for β-catenin (Chen et al., 2002).

The mobility of all tested proteins significantly decreased during the maturation from a calcium-dependent to the calcium-independent switch to the hyper-adhesion state. The already low mobile fractions almost halved for the DCs, PG and DP, and decreased by 15% for Pkp2a. A similar result was found recently when hyper-adhesion was pharmacologically induced with Gö6976, an inhibitor of conventional PKC isoforms, though Pkp was not included in that study (Bartle et al., 2020). The disadvantage of using such an inhibitor is that it may block multiple cellular events, any one of which may directly or indirectly affect the mobility of desmosomal components. Now that we have studied simple time-dependent maturation of desmosomes, we may be fully confident that stabilisation of desmosomal components is indeed a consequence of hyper-adhesion.

Of particular interest is the decrease in mobility of DP that accompanies desmosome maturation. During the calcium-dependent phase we found DP to be substantially more mobile than the stable core represented by the DCs and PG, but with the onset of hyper-adhesion its mobility was reduced to the extent that it could now also be considered part of the core. We suggest that this stabilisation of DP makes a significant contribution to the gaining of hyper-adhesion. This view is consistent with data showing that a phospho-null DP mutant with enhanced keratin binding properties leads to an increased association of the desmosomal plaque with keratin filaments, resulting in decreased protein exchange and increased desmosome stability (Albrecht et al., 2015; Bartle et al., 2020). It was proposed that inhibition of PKC inhibitors or downregulation of PKCα may promote DP-keratin integration leading to the onset of hyper-adhesion (Bartle et al., 2020). Furthermore, during formation of the desmosome-keratin scaffold the development of radial and inter-desmosomal keratin filaments coincided with a decrease in DP mobility (Moch et al., 2019).

There is also evidence for alternative ways of influencing the mobility of desmosomal proteins, one deriving from crosstalk with the actin cytoskeleton. For example, the knockdown of α-adducin, a protein involved in the assembly of a plasma membrane-stabilising cortical spectrin-actin network, reduces Dsg3 mobility in the membrane (Hiermaier et al., 2019). Adducin is a PKC substrate (Larsson, 2006), as are many of the actin regulatory proteins. Also, actin-mediated endocytosis is regulated by PKCα and it can be blocked by the inhibitor Gö6976 (Hryciw et al., 2005), as can the actin-mediated internalisation of desmosomes (Holm et al., 1993). Furthermore, it has been shown that complete absence of PKCα in mice appears to stabilise hyper-adhesion of desmosomes in epidermal wounds (Thomason et al., 2012). It is thus highly probable that PKCα regulates the mobility and the life cycle of the desmosomal complex at multiple levels, even though it is not required for desmosome assembly (Thomason et al., 2012). Further experiments are needed to determine the precise role of PKCα in the life cycle of desmosomes.

Another possible reason for such decrease in mobility of desmosomal components could be the progressive packing of cadherins into a higher order array that locks DCs in a caclium-independent adhesion state; this increasing order undoubtedly occurs during the onset of hyper-adhesion (Garrod et al., 2005; Kimura et al., 2012). This suggestion was disputed by Bartels and colleagues who found that the ordered packing of Dsg3 with a genetically modified extracellular domain was lost on chelation of Ca^2+^ from pharmacologically induced hyper-adhesive desmosomes (Bartle et al., 2020). However, the modified Dsg3 may not reflect the behaviour of wild type DCs, since it is difficult to understand how adhesion between opposed desmosomal halves can be maintained in the presence of chelating agents, the defining criterion of hyper-adhesion, if the ordered array of their EC domains is lost (Garrod, 2013).

Irrespective of their maturation state, desmosomes form extremely strong intercellular bonds. Our data demonstrate that cells appear to move apart by tearing part of the desmosome-IF complex out from the neighbouring cell, resulting in the internalisation of the desmosomes as a whole unit with associated keratin filaments. This observation was surprising as others claimed that cell scattering involved desmosome dissociation or junctional splitting together with partial desmosome disassembly (Boyer et al., 1989; Savagner et al., 1997). We found no evidence for such mechanisms of desmosome dismantling in cells undergoing scattering. The forceful internalisation of the whole complex by one of the attached cells recalls the observation of whole desmosome internalisation seen in vivo in wound edge keratinocytes and certain carcinomas (Allen and Potten, 1975; Garrod et al., 2005). It is striking that once internalised, desmosomes remained intact as judged by both their ultrastructure and protein composition. Over longer periods after internalisation we previously showed that components of engulfed desmosomes remained associated and were not recycled, but rather underwent lysosomal and proteasomal degradation (McHarg et al., 2014). We note that a similar mode of internalisation to that we describe here has also been reported for tight junctions (Matsuda et al., 2004).

In their study of scattering by NBT II cells, Boyer et al. noted that loss of cell-cell contact was accompanied by the appearance within the cytoplasm of dots containing desmosomal components and associated keratin filaments (Boyer et al., 1989). We suggest that these dots were equivalent to the whole internalised desmosomes that we report here. We have further shown that some of these filaments are contributed by the opposing cell of an adjacent pair, indicating that the filaments have been fragmented during cell separation. If we take the view that cell separation occurs by tearing under the force generated by cell movement, this would suggest that the keratin filaments have simply broken under the applied force. However keratins are the strongest filament component in the cell, with particular high tensile strength. Alternatively, we may speculate that a signalling event such as phosphorylation is involved in severing the keratin filaments and that such a signalling process would therefore have a facilitating role in cell separation. This would be analogous to the rapid and specific phosphorylation events that sever cytoskeleton components for daughter cell separation at the end of cytokinesis (Goto et al., 2000).

In conclusion, our data represented here leads to a new concept of dualistic desmosome organisation. There appear to be two protein modules or fractions within the junctions, i.e. a stable core which is presumably responsible for the strong adhesive function of the desmosome, and a much more mobile protein fraction, exemplified by Pkp2a, which has the potential for signalling and regulatory function. This behavioural dichotomy is both present and detectable when desmosomes are relatively newly formed and calcium-dependent, and when they mature to calcium-independence. This dualism persists even when desmosomes are being down-regulated and removed.

## ACKNOWLEDGEMENTS

This work was supported by the Faculty of Biology, Medicine and Health at the University of Manchester, UK and by the Agency for Science, Technology and Research (A*STAR), Singapore. We thank the staff of the Bioimaging facility at the University of Manchester and the A*STAR Microscopy Platform for their help with imaging and analysis. C. Ballestrem acknowledges the Biotechnology and Biological Sciences Research Council (BBSRC) and the Wellcome Trust for funding of this project. The C. Ballestrem laboratory is part of the Wellcome Trust Centre for Cell-Matrix Research, University of Manchester, which is supported by core funding from the Wellcome Trust (grant number 203128/Z/16/Z). The work in Singapore was supported by institutional core funding to the Skin Research Institute of Singapore from the Biomedical Research Council of Singapore. J. Fülle is supported by the Faculty of Biology, Medicine and Health at the University of Manchester and by the Agency of Science Technology and Research (A*STAR).

## AUTHOR CONTRIBUTIONS

JBF, HH, DG and CB designed experiments, with input from EBL and GDW. JBF, HH, DL and WIG performed the experiments and analysed the data. RA and BY provided assistance with generation of tagged desmosomal constructs. Figures were prepared by JBF. JBF, DG, EBL and CB wrote the manuscript. CB, DG, GDW and EBL supervised the project and acquired funding.

## Materials and Methods

### Cell lines and transfection

Madin-Darby canine kidney II cells (MDCK) (Madin and Darby, 1958) (ECACC) were cultured at 37°C in 5% humidified CO2 in high glucose Dulbecco’s Modified Eagle Medium (DMEM, Sigma or GE Healthcare), supplemented with 4 mM L-glutamine, 10% (v/v) foetal calf serum (FCS, Gibco or GE Healthcare) and 100 U/ml penicillin and 100 µg/ml streptomycin (P/S; Gibco), herein referred to as standard medium (SM) with a calcium concentration of approximately 1.8 mM. Cells were transfected using Lipofectamine 3000 transfection reagent, according to the manufacturer’s instructions (Invitrogen). MDCK cells stably expressing Dsc2a-YFP were kindly provided by R.E.Leube, University Hospital RWTH, Aachen (Windoffer et al., 2002). For live-cell imaging experiments, the cells were plated on glass-bottom dishes (MatTek or Ibidi). For cell scattering experiments using HGF, the glass-bottom dishes were coated with 10 µg/ml bovine plasma fibronectin (FN; Sigma).

### Cloning and constructs

DP-EGFP (32227), Dsg2-mCherry (36991) and mCherry-KRT18 (55065) were purchased from Addgene. mEmerald-E-cadherin was obtained from the Michael Davidson collection 54072 from P. Kanchanawong (Mechanobiology Institute, Singapore). Dsc2a-YFP was a kind gift from R. E. Leube. To generate constructs containing PaGFP, neonGreen and mScarlet, we have used plasmids where the desmosome genes were already cloned into a custom-made vector by Oxford Genetics where the vector pSF (OG394R1) was modified to have an EF1a promoter and puromycin selection marker. The vector was linearized by restriction digestion followed by gel purification (see table 2 for restriction enzymes used), and the purified vector was used to clone fragments containing the ORFs of AGFP, neon-Green and mScarlet, these fragments were amplified by Polymerase Chain Reaction using Phusion® High-Fidelity DNA Polymerase (M0530L, New England Biolabs (NEB)) using 35 cycles with an annealing temperature of 60 °C (see table 2 for list of primers) and cloned by Gibson Assembly Cloning Kit (EE5510S, NEB). All primers were designed using SnapGene (GSL Biotech LLC, Chicago, IL) and were synthesised by Eurofins Genomics (Germany). All ORF sequences were confirmed by Sanger sequencing by Eurofins.

### Antibodies and reagents

Primary antibodies and dilutions used were: anti-desmoplakin I and II (clone 11-5F, D. R. Garrod), 1:400; anti-plakoglobin (clone 15F11, Sigma), 1:100; anti-desmocollin 2 and 3 (clone 7G6, Thermofisher), 1:100; anti-keratin 8 (clone LE41, E. B. Lane), used as hybridoma culture supernatant; anti-plakophilin 1 (PP1-5C2, Abnova), 1:50. Secondary antibodies conjugated to Alexa Fluor 488, 594 or 647 were all from Thermofisher (used at 1:500). Alexa Fluor 488- (1:500), Texas-Red-X- (1:500) and Alexa Fluor 647-conjugated (1:200) phalloidin were from Life Technologies.

### Calcium switch assay

MDCK cells were cultured in SM. They were washed thrice with calcium- and magnesium-free PBS (Gibco) and incubated with calcium-free DMEM (Gibco) supplemented with 10% chelated FBS and 3 mM EGTA (Sigma-Aldrich), herein referred to as “+EGTA” for 1 h (unless specified otherwise) at 37°C 5% humidified CO2. Calcium sensitivity of desmosomes was quantified as previously described (Wallis et al., 2000). In brief, following Ca^2+^-chelation, the cells were fixed in ice-cold methanol for 10 min and stained for desmoplakin by immunofluorescence. Cells which remained attached by desmoplakin positive projections were scored as having calcium-independent desmosomes and expressed proportional to the total cell number.

### Microscopy

Co-localisation experiments were carried out using a Delta Vision microscope (Applied Precision) 60x/1.42 Plan Apo N (Oil) objective and a Sedat Quad filter set, with images collected using a Retiga R6 (Q-Imaging) camera. 3D-SIM was performed using a DeltaVision OMX Version 4 Blaze microscope (GE Healthcare), equipped with 405-, 488-, and 568-nm lasers and a BGR filter drawer. A 100x/1.40 PSF Plan Apo Oil objective and liquid-cooled Photometrics Evolve EM charge-coupled device camera for each channel were used. An ImageJ macro was used for post-acquisition alignment. FLAP and FRAP experimental data was collected were acquired using an inverted Nikon Eclipse Ti microscope equipped with a 100×/NA 1.40 Plan Apo oil immersion objective lens, a focus drift correction system, a piezo-motorized stage, a 37 °C on-stage incubation system (LCI), 100 mW diode lasers (405 nm, 491 nm and 561 nm), an EMCCD camera (Evolve 512; Photometrics), CSU-22 spinning disk scan head (Yokogawa) and a 3D FRAP system (iLAS2 Roper Scientific). The microscope system was controlled using MetaMorph (Molecular Devices) and iLAS2 (Roper Scientific) software.

### Immunofluorescence microscopy

Cells were fixed with 4% (w/v) paraformaldehyde in PBS or 100% ice-cold Methanol for anti-desmoplakin (clone 11-5F) immunostaining. Antibodies were diluted in 1% BSA (Sigma-Aldrich) and added to the cells for 1h. Images were acquired on the Delta Vision systems (above) and processed using the FIJI-ImageJ software (version 1.53g). To analyse xy, background was subtracted using a rolling ball algorithm. Pearson’s correlation was calculated using the Coloc 2 plugin of Fiji. RGB profiler was used to generate intensity profiles.

### Live-cell imaging

#### FRAP

MDCK cells expressing neonGreen, mEmerald or EGFP constructs were seeded at confluent density (1.35 × 105 cells/cm2) 3 days and 24 hours before imaging on uncoated glass bottom dishes in SM. Images were acquired using the Nikon Eclipse Ti system (above). For FRAP quantification five ROIs of 2 µm diameter circles at the cell-cell junctions were selected and photobleached with a 10 ms burst of the 488 nm laser at 100% power. iLAS2 software was used to capture three images prior to photobleaching and one image every 10 s for 5 min to 10 min post bleaching. Movies were analysed using Fiji (version 1.53g) by manually tracking the ROI and measuring the fluorescence signal. The subsequent analysis was performed as described previously (Carisey et al., 2011).

#### FLAP

MDCK cells co-expressing the required PaGFP-tagged constructs and an mScarlet-tagged desmosome marker protein (Dsc2a-mScarlet unless otherwise specified) were plated at confluent density (1.35 × 105 cells/cm2) 3 days and 24 hours before imaging on uncoated glass bottom dishes in SM. Images were acquired using the Nikon Eclipse Ti system (above). Similarly to FRAP five ROIs of 2 µm diameter circles were selected and photoactivated using a 405 nm laser at 100% power for 10 ms. iLAS2 software was used to capture three images prior and one image every 10 s for 5 min up to 1 h post photoactivation. Movies were analysed using Fiji (version 1.53g) and quantified as described previously (Atherton et al., 2015).

#### CLEM

MDCK cells stably expressing Dsc2a-YFP were co-cultured with MDCK cells transiently expressing Dsg2-mCherry on a gridded glass-bottom dish (MatTek). Cells were treated with HGF (see above). Live-cell imaging of regions of interest (ROI) was carried out using the Nikon Ti Eclipse microscope (see above). Cells were grown on glass-bottom dish with gridded coverslip (MatTek Corporation, Part No: P35G-1.5-14-CGRD). After fluorescence images of cells co-expressing Dsc2a and Dsg2 were acquired, bright field (phase contrast) images of cells and finder grid index were captured at lower magnification for ROI reference. Cells in dishes were then fixed with 2% glutaraldehyde and 4% formaldehyde in 0.1M Cacodylate buffer, post-fixed in 1% Osmium tetroxide, dehydrated in ethanol and embedded in Epoxy resin (EPON 812, SERVA). Upon resin polymerization the coverslip was removed using liquid nitrogen (leaving finder-grid embossed to the block surface) and respective ROI were re-localized under stereomicroscope. Target cells within ROI were cut horizontally with ultramicrotome (LEICA Ultracut-UC7), ultrathin sections collected on formvar/carbon-coated slot grids (Ted Pella Inc., 01805-F) and post-contrasted with Uranyl Acetate and Lead Citrate. Sections were analysed under JEOL JEM-1010 transmission electron microscope operated at 80kV and images acquired with SIA 12C CCD camera. Desmosome width was measured by taking the average of three line measurements per desmosome using ImageJ. 10 desmosomes at the membrane and following internalisation were measured.

### Statistical analysis

Graphing and statistical analysis were performed by using GraphPad Prism 9.0.0 software. When comparing means, the D’Agostino–Pearson test was used to assess the normality of the data to determine the appropriate statistical tests to use.

---

**Supplemental Fig. 1.**
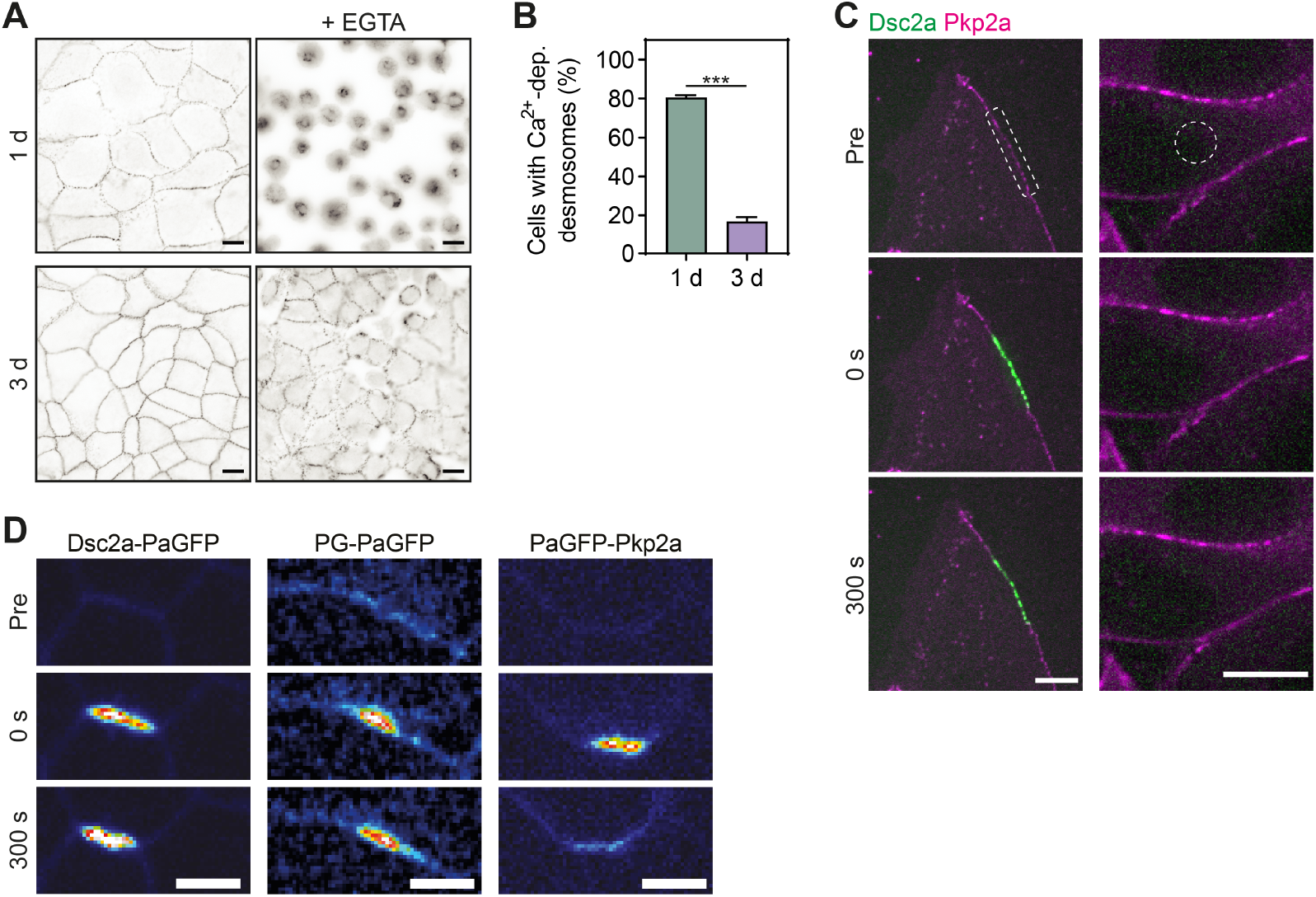
Differential desmosomal protein dynamics depends spatial and temporal conditions. **A**. Representative images of wt MDCK cells acquiring calcium-independence. The cells were cultured confluent for 1 or 3 d, treated Ca^2+^-chelating medium (+ EGTA) and stained for desmoplakin. Scale bar: 10 µm.**B** Quantification of transfected MDCK cells with calcium-dependent desmosomes (1d N=69; 3d N=46).**C** Time-lapse images of FLAP of 1 d cultured MDCK cells transfected with Dsc2a-PaGFP and mScarlet-Pkp2a comparing the photoactivation at the membrane and cytoplasmic activation. Note to absence of a cytoplasmic pool of Dsc2a. Scale bars: 10 µm.**D** Time-lapse images of FLAP comparing Dsc2a-PaGFP, PG-PaGFP and PaGFP-Pkp2a in MDCK cells cultured for 3 d. Images are presented in a colour intensity scale. Scale bars: 5 µm.

**Supplemental Fig. 2.**
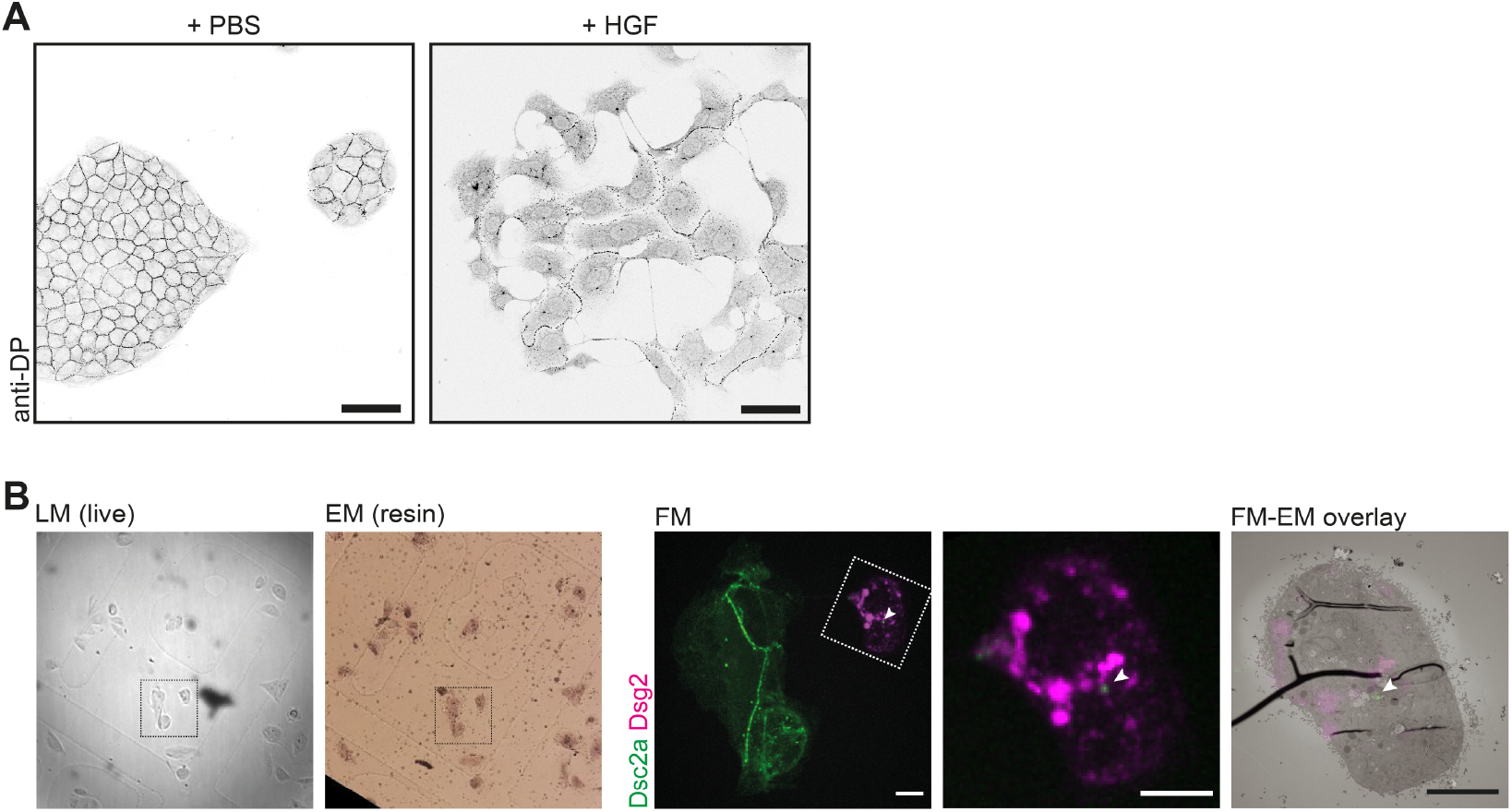
Hepatocyte growth factor-induced cells scattering leads to whole desmosome internalisation. **A** MDCK cells were cultured sub-confluent for 2 d, serum-starved for 12 h, treated for 4 h with 40 ng/ml hepatocyte growth factor (HGF) or same v/v PBS, and stained for desmoplakin (DP). Scale bars: 50 µm. **B** In same way treated mixed populations of MDCK cells expressing either Dsc2a-YFP (green) or Dsg2-mCherry (magenta) were used for correlative light and electron microscopy to identify cells which had internalised double-labelled desmosomes. LM = light microscopy, EM = electron microscopy, FM = fluorescent microscopy. Scale bars: 10 µm.

